# Peripheral tolerance checkpoints imposed by ubiquitous antigen expression limit antigen-specific B-cell responses under strongly immunogenic conditions

**DOI:** 10.1101/2020.03.10.985127

**Authors:** Jeremy F. Brooks, Peter R. Murphy, James E.M. Barber, James W. Wells, Raymond J. Steptoe

**Author notes:** Corresponding author: R. J. Steptoe, UQ Diamantina Institute, Level 6, Translational Research Institute, 37 Kent Street, Woolloongabba, Queensland, Australia 4102. Author contributions: JFB and RJS designed and performed experiments, analysed data and wrote the manuscript. PRM, JEMB performed experiments. JWW provided intellectual insights and edited the manuscript.

## Abstract

A series of layered peripheral checkpoints maintain self-reactive B cells in an unresponsive state. Autoantibody production occurs when these checkpoints are breached, however, when and how this occurs is largely unknown. In particular, how self-reactive B cells are restrained during bystander inflammation in otherwise healthy individuals is poorly understood. A weakness has been the unavailability of methods capable of dissecting physiologically-relevant B-cell responses, without the use of an engineered B-cell receptor. Resolving this will provide insights that decipher how this process goes awry during autoimmunity or could be exploited for therapy. Here we use a strong adjuvant to provide bystander innate and adaptive signals that promote B-cell responsiveness, in conjunction with newly developed B cell detection tools to study in detail the ways that peripheral tolerance mechanisms limit the expansion and function of self-reactive B cells activated under these conditions. We show that although autoreactive B cells are recruited into the germinal centre, their development does not proceed, possibly through rapid counter-selection. Consequently, differentiation of plasma cells is blunted, and autoantibody responses are transient and devoid of affinity maturation. We propose this approach and these tools can be more widely applied to track antigen-specific B cell responses to more disease relevant antigens, without the need for BCR transgenic mice, in settings where tolerance pathways are compromised or have been genetically manipulated to drive stronger insights into the biology underlying B cell-mediated autoimmunity.

## Introduction

Central B-cell tolerance mediated by clonal deletion and receptor editing in the bone marrow is incomplete and self-reactive B cells escape into the peripheral repertoire. To counter this, many overlapping and complementary peripheral tolerance mechanisms have evolved to limit the responsiveness of potentially self-reactive B cells. Peripheral B-cell tolerance normally arises when B cells encounter cognate antigen in a way that leads to ‘unproductive’ B-cell receptor (BCR) signaling with chronic antigen recognition being most potent, and likely essential for this. Chronic antigen stimulation leads to B cell receptor (BCR) desensitization and anergy (1, 2). Anergic B cells are unable to effectively solicit T-cell help(3), are excluded from entry into follicles (4, 5), have a curtailed lifespan (4) and are deleted from the recirculating repertoire. Cell surface and membrane-bound self-antigens, which are BCR-accessible, are potent inducers of B-cell peripheral tolerance (6, 7) whereas soluble proteins are less effective (8). Aside from these B-cell intrinsic mechanisms, the presence or absence of cognate antigen-specific T-cell help is a major controlling influence on the productivity of B-cell responses. Because strong T-cell tolerance is present for soluble self-proteins (6, 9), B-cell responses and autoantibody production to soluble self-proteins may be limited primarily by loss of T-cell help (9).

Despite the many mechanisms that control self-reactive B cells, in autoimmune diseases, central and/or peripheral tolerance mechanisms fail. Transcriptional and molecular analyses have revealed many pathways that control B-cell responsiveness (10) and genetic studies have defined where control pathways or mechanisms can fail in autoimmune diseases (reviewed in (11). Other studies identify ‘accessory signals’ capable of breaking peripheral B-cell tolerance or bypassing the checkpoints that normally control B-cell responsiveness including innate signalling (TLRs), T-cell collaboration (CD40L) and dendritic cell input (BAFF) (12). Some of these mechanisms are of relevance to autoimmune disease (13) and provide valuable insights into how autoimmune disease is initiated or perpetuated. Importantly, understanding how the responsiveness of self-reactive B cells is controlled will provide important insights into how B-cell tolerance can be engineered for therapeutic goals. This would be particularly pertinent for approaches such as gene therapy where antigens of interest, to which tolerance is desired, are expressed chronically or ubiquitously enabling normal B-cell tolerance mechanisms to be exploited.

The limited availability of tools to accurately monitor naïve endogenous self-reactive B cells in non-BCR-engineered models has been a challenge to progressing the understanding of B-cell tolerance. Most insights have been gained in models where the BCR repertoire is genetically manipulated, typically to increase the clonal frequency, or for adoptive transfer, to enable antigen-specific B cells to be detected. However, this has the consequence of altering tolerance outcomes (14). We aim to understand the robustness of peripheral tolerance and B-cell checkpoints in a setting devoid of BCR manipulation and where a ‘self-antigen’ is expressed ubiquitously at the cell surface. A key goal is to understand whether the knowledge gained using BCR-Tg settings also stands for setting where there is no genetic manipulation of BCR. We employ newly-developed tools to dissect the fate and function of self-reactive B cells after administration of antigen with a potent adjuvant. In detailed analyses we found, despite the provision of strong adjuvant signals, self-reactive B-cell responses were transient, affinity maturation was restrained and long-lived plasma cell and memory B cells (Bmem) formation was curtailed, thus demonstrating the robustness of peripheral B-cell checkpoints under these conditions in a non-autoimmune background. The system explored here has revealed meaningful insights into control of B-cell responsiveness under ‘normal’ non-BCR engineered conditions and would form the basis of a useful system in which to test the influence of genetic and environmental factors on the outcome of peripheral B-cell tolerance.

## Materials and Methods

### Mice

Actin.mOVA (act.OVA) mice ubiquitously expressing membrane-bound ovalbumin (OVA) under the control of a β-actin promoter (15) were maintained at the TRI Biological Resources Facility (Brisbane, Australia) by heterozygous backcrossing to C57BL/6J/Arc (Animal Resources Centre, Perth Australia). Male and female, age-matched (typically within 2-4 weeks) transgenic and non-transgenic littermate 6-16 weeks of age were used. Studies were approved by the University of Queensland Ethics Committee.

### Immunizations

OVA (Sigma, Grave V, A5503) 2mg/mL in PBS was emulsified in Complete Freund’s Adjuvant (CFA, Sigma F5881 or Difco 23810, MO USA) or Incomplete Freund’s Adjuvant (IFA, Sigma F5506, MO USA) using a mini-BeadBeater (Biospec Products, OK-USA) and 100μl injected s.c. at the tail base. In some experiments, 100μg OVA was administered i.p. with 50μg anti CD40 (FGK-45, BioXCell – NH, USA) on day 0, with additional anti-CD40 (i.p.) at day 2, 4, 6, 8. In some experiments, 50μg OVA was mixed with 20μg Quil-A (Sigma) in PBS and injected s.c. at the tail base. For adjuvant-free conditions, OVA was either injected alone or depleted of LPS as described (16) and 100μg injected s.c. at the tail base. Where sham-immunized controls were included, PBS and adjuvant was prepared and injected as described for OVA immunizations.

### Flow cytometry

OVA-labelled tetramer preparation has been described (17). OVA and hen egg lysozyme (HEL) were biotinylated in-house and tetramerised with streptavidin-PE (Biolegend, CA USA or Prozyme, -- CA, USA), streptavidin-APC (Biolegend, CA USA) or streptavidin-BV510 (BD Bioscience, CA USA) for two hours on ice. OVA tetramers were used at 14.6 - 21.2nM (depending on batch) for staining. HEL tetramers were used at 6.6 - 7.3nM. IgM-PECy7 (RMM-1), IgD-BUV395 (11-26c.2a), CD4-FITC (GK1.5), CD8-FITC (53-6.7), CD11c-FITC (N418), B220-APC (RA3-B62), CD19-APC (6D5) or CD19-BV785 (6D5) and GL7-Pacific Blue (GL7) were purchased from Biolegend (CA USA). CD138-AF700 (281-2), GL7-BV421 (GL7), IgG1-biotin (A85-1), and IgG2a,2b-biotin (R2-40) were purchased from BD Bioscience (CA USA). NK1.1 (PK136), Gr1 (RB6-8C5) and CD11b (M1-70) were purchased from BioXCell (NH USA) or grown in-house and labelled with FITC in-house.

Draining lymph nodes (inguinal) from each immunized mouse were collected, pooled and pressed through a 70μm strainer (Corning, NY, USA) to create single-cell suspensions. For analysis of bone marrow, femurs and tibiae from immunized mice were harvested and BM flushed with cold PBS/FCS (2.5% v/v). BM was repeatedly passaged through a 21g needle to prepare a single cell suspension and erythrocytes lysed using ammonium chloride buffer. Antibody and tetramer staining was as described (17) and all staining steps were performed on ice. For intracellular tetramer staining, cell suspensions were stained for CD138-AF700, CD19-BV785 and FITC-conjugated lineage mix. In some experiments, cells were also stained with Sca-1 PerCP-Cy5.5 (Biolegend, CA USA). Cells were then fixed (Fixation buffer, #420801, Biolegend, CA USA) and permeabilized (Fix/Perm buffer, #55473, BD Pharmingen, CA USA) and stained with tetramer (OVA-PE tetramer only) as described for surface tetramer staining (17).

Absolute cell number was enumerated using Flow-Count™ fluorospheres (Beckman Coulter, CA USA) as described (18). Data were collected on an LSR Fortessa X-20 (BD Bioscience, CA USA) and analysed using Diva Version 8 (BD Bioscience, CA USA).

### ELISA

For OVA-specific IgG1, IgG2b, IgG2c or IgM titres, plasma was serially diluted (4-fold dilutions) and added to ELISA plates (Maxisorp, Nunc, MA USA) pre-coated with 10μg/ml OVA (Sigma Grade V, A5503 MO USA). OVA-specific antibody was detected using biotinylated anti-mouse antibodies (anti-IgG1 [RMG1-1] 0.02μg/ml, anti-IgG2b [RMG2b-1] 0.05μg/ml, Biolegend, CA, USA; anti-IgG2c 0.2μg/ml, Southern Biotech, CA USA; anti-IgM 0.2μgml, Thermo Fisher MA USA) followed by SA-HRP (1:2000 dilution, #P030701-2, Dako, CA USA), and visualised with TMB (#421101, Biolegend, CA USA). Absorbance was measured at 450nm and corrected for absorbance at 540nm using a MultiSkan Go spectrophotometer (Thermofisher, MA USA).

### Affinity ELISA

The affinity ELISA was adapted from a previously published method (19). Absolute OVA-specific IgG1 concentration was determined for each sample by ELISA as described above using OVA-14 (Sigma, MO USA) as a standard and equalized across all samples. Samples were blocked with increasing concentrations of OVA (10^−4^ to 10^−12^M), left unblocked or blocked with an irrelevant antigen (HEL, 10^−4^M). Samples were then transferred to ELISA plates pre-coated with OVA (1μg/ml). The assay then proceeded as described above for OVA-specific IgG1 ELISA.

### Statistical analysis

Comparison of means used student t-test or ANOVA with Tukey post-hoc analysis for multiple comparisons for normally distributed data and Kruskal-Wallis and Mann-Whitney test for non-normally distributed data. (GraphPad Prism V8, GraphPad Software, Inc., CA USA). For antibody titres, statistical testing was performed on log-transformed data. Correlations were performed using a semi-log linear regression (y-log axis, x-linear axis) or log-log regression (GraphPad Prism V8, GraphPad Software, Inc., CA USA).

## Results

### Provision of highly immunogenic stimuli partially overrides peripheral B-cell tolerance checkpoints

Our aim was to understand how peripheral checkpoints might control the response of B cells to immunogenic stimuli in mice that are rendered ‘tolerant’ through constitutive expression of a model self-antigen. We first tested the outcome of immunization of non-Tg controls and ovalbumin (OVA)-expressing act.OVA mice with OVA and a range of adjuvants that might carry diverse or different accessory signals (**Fig. 1A**). For this we used OVA or OVA that had been treated to remove LPS (LPS-free OVA), each without any adjuvant, OVA emulsified in incomplete Freund’s adjuvant (OVA/IFA) or complete Freund’s adjuvant (CFA, OVA/CFA). We also immunized with OVA in conjunction with serial injections of agonistic anti-CD40 mAb or OVA with Quil-A. In non-Tg controls there was a graded response to immunization 14 days after immunization. OVA without adjuvant induced modest OVA-specific IgG1 levels and, notably, OVA-specific IgG1 production was abrogated by LPS depletion. OVA/CFA and OVA/IFA, however, led to the production of substantial levels OVA-specific IgG1 in non-Tg controls, (**Fig. 1B**). Other immunization regimes led to intermediate levels of OVA-specific IgG1 production, similar to OVA without adjuvant. In contrast to non-Tg controls, in act.OVA mice only OVA/CFA reliably led to production of detectable levels of OVA-specific IgG1 and the overall level was very low (**Fig 1B**, ∼0.2ug/ml). Therefore, a strong adjuvant was required to elicit OVA-specific antibody production in OVA-expressing mice and even then, minimal amounts of OVA-specific antibody were produced. Interestingly, in act.OVA mice, both OVA and LPS-free OVA without adjuvant were incapable of eliciting OVA-specific IgG1, suggesting that OVA-specific B cells in act.OVA were largely unresponsive and this manifests here as an insensitivity to the adjuvant effects of endotoxin relative to non-Tg mice.

**Figure 1.**
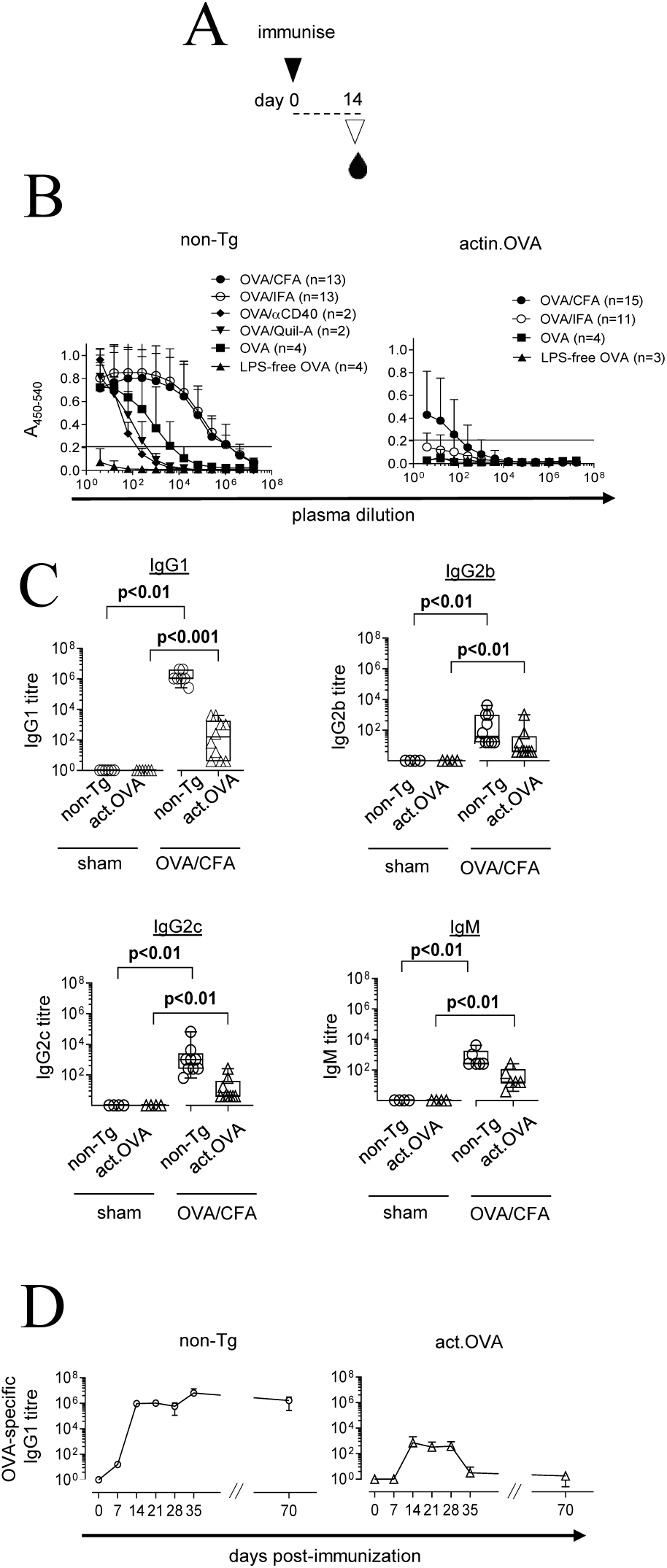
OVA/CFA partially overrides peripheral B-cell checkpoints. A) Schematic of experimental design. Non-Tg and act.OVA mice were sham-immunized (PBS/CFA) or immunized with OVA/CFA, OVA/IFA, OVA/antiCD40, OVA/Quil-A or OVA without adjuvant and then OVA-specific IgG1 ELISAs performed on plasma collected 14 days later (B,C) or at the timepoints indicated (D). B) Titration curves for non-Tg and act.OVA mice immunized as indicated. C) OVA-specific IgG1, IgG2b, IgG2c and IgM titres for OVA/CFA immunized and sham-immunized mice. D) Time-course of plasma OVA-specific IgG1 titres determined at the indicated timepoints. Data show mean±SD with number of replicates indicated (B), individual values pooled from 2-6 experiments with median and interquartile range shown (C) or mean±SD (n=3-10). Mann-Whitney test (C) or ANOVA with Tukey’s post-test (D) of log-transformed data

When antibody titres for IgM, IgG1, IgG2b and IgG2c were plotted for individual mice immunized with OVA/CFA, non-Tg mice produced anti-OVA antibodies of all isotypes (**Fig. 1C**). Notably act.OVA mice produced significantly lower amounts of OVA-specific antibodies across all isotypes tested (**Fig. 1C**). Therefore, the strong adjuvant CFA provides immunostimulatory signals that partially overcome peripheral B-cell checkpoints, potentially through diverse innate signals or from bystander T-cell responses activated by the mycobacterial (Mtb) components of CFA, which are absent from the similar, but Mtb-free, IFA.

To gain more insight into the nature of the antibody response induced by OVA/CFA immunization we monitored OVA-specific IgG1 in the circulation. In non-Tg controls, OVA-specific IgG1 titres rose rapidly (detectable by day 7), reached a maximum 14 days after immunization and then remained stable at this level until at least day 70 after immunization (**Fig. 1D**). By contrast, in act.OVA mice, OVA-specific IgG1 accumulation was slower (not detectable at day 7), was much lower overall and accumulation was transient peaking 14 days after immunization and diminishing to near undetectable levels by day 35 after immunization (**Fig. 1D**).

### OVA-specific B cell activation and germinal centre differentiation is limited when OVA is widely expressed

To understand in more detail which specific components of responsiveness to OVA were modulated by widespread OVA expression we analysed B-cell responses at the cellular level (**Suppl Figure 1, Fig 2A**). After immunization, the overall B cell population was expanded in both non-Tg and act.OVA mice to a similar extent relative to naïve controls (**Supplementary Fig. 2A**) and germinal centre (GC) B cell differentiation was present in sham- (PBS/CFA) and OVA/CFA-immunized non-Tg and act.OVA mice although the scale may have been larger in OVA/CFA immunized non-Tg mice (**Fig. 2B-D**).

**Figure 2.**
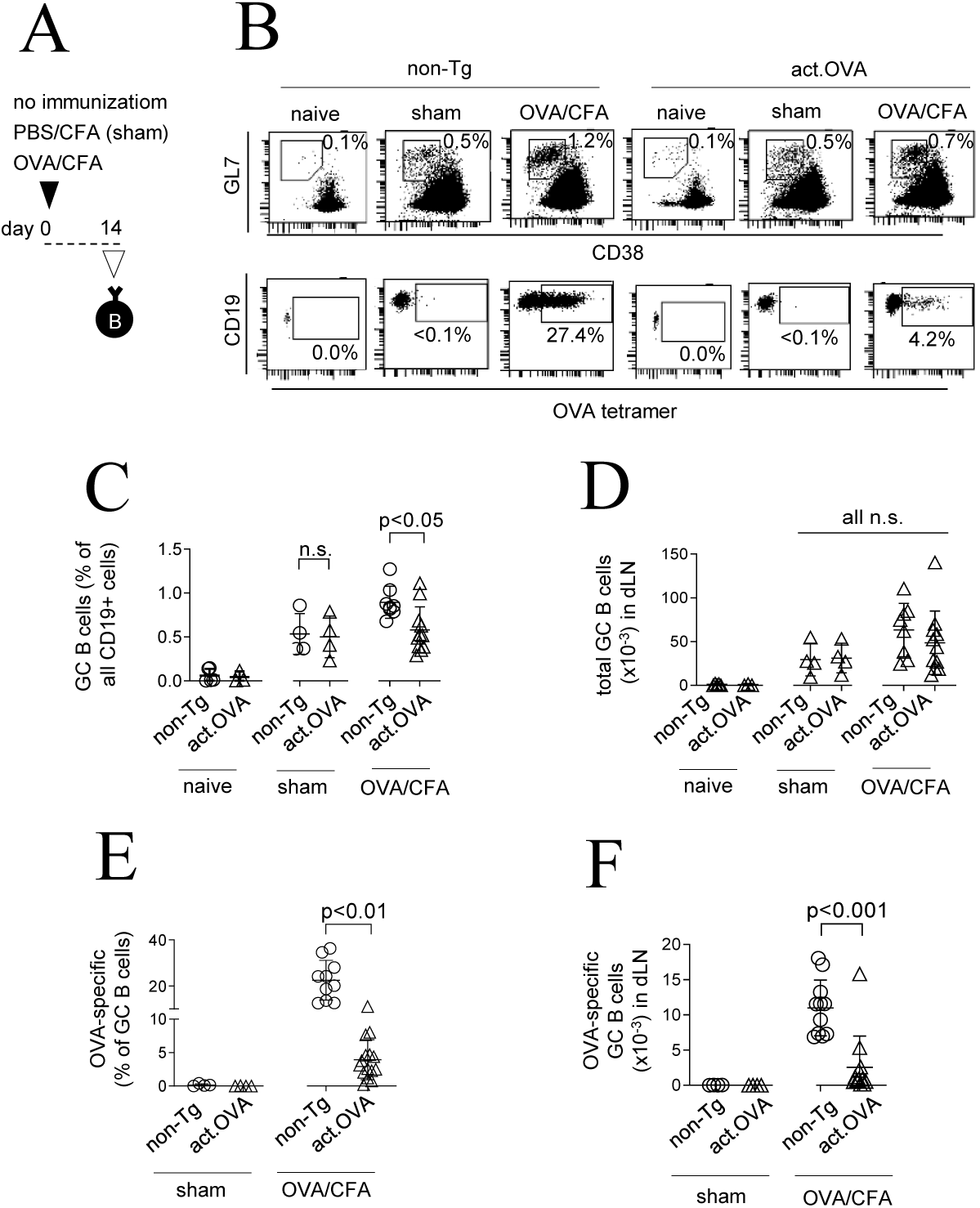
OVA-specific GC responses are blunted in act.OVA mice. A) Schematic of experimental design. B) Representative FACS plots showing gating for GC B cells within lin^-^ CD19^+^ B cells in pooled draining (inguinal) lymph nodes (dLN) (top), and gating for OVA tetramer binding (bottom). C) Total GC B cell frequency as a proportion of all CD19^+^ B cells. D) Total GC B cell number per dLN. E) OVA-specific GC B cells as a proportion of all GC B cells. F) total number of OVA-specific GC B cells per dLN. Data show representative FACS plots (B) or values for individual mice with mean ± SD (C-F). Data are pooled from 3-5 experiments. ANOVA with Tukey’s post-test (C-F).

Given the responsiveness observed in both sham- and OVA/CFA immunized act.OVA mice, which may be directed at MTb antigens in CFA, we next used tetramers to dissect the OVA-specific components of the B-cell response to immunization. Based on antibody production (**Fig. 1**), we speculated that OVA-specific GC B-cell differentiation may have occurred in act.OVA mice despite widespread expression of OVA. In non-Tg mice, a large proportion (∼25%) of GC B cells were OVA-specific cells, whereas they formed a significantly smaller fraction (∼5%) in act.OVA mice (**Fig. 2E**) such that OVA-specific GC B cells were ∼4.3-fold more numerous in non-Tg mice than in act.OVA mice after OVA/CFA immunization (**Fig. 2F**). Overall, failure of OVA-specific GC B cells to develop in sham-immunized mice and their reduced proportion among total GC B cells in act.OVA mice supports the premise that a large component of the GC response in both non-Tg and act.OVA mice is to the result of adjuvant-associated signals (or antigens). Nevertheless, the presence of OVA-specific B cells participating in ongoing GC responses in act.OVA mice confirms a partial breach in peripheral tolerance in the presence of immunogenic adjuvant possibly as a consequence of bystander T-cell help provided through adjuvant-associated signals.

### Affinity maturation but not class-switch recombination of OVA-specific GC B cells is impaired in act.OVA mice

When self-reactive B cells are inappropriately recruited into a GC, affinity counterselection regulates the GC-plasma cell (PC) axis to prevent the ultimate production of self-reactive antibodies (20). Surprisingly, the extent of OVA-specific GC development was quite substantial after OVA/CFA immunization of act.OVA mice. Therefore, we sought to identify whether the typical outcomes of a GC reaction such as generation of isotype-switched, affinity matured and long-lived antibody responses occurred. Analysis of IgG^+^ (IgG1, IgG2b), class-switched B cells revealed their total number was relatively similar for OVA/CFA-immunized non-Tg and act.OVA mice (**Supplementary Figure 2B**). However, the frequency of OVA-specific cells within the switched B cell population (**Fig. 3A**) was remarkably similar to that for OVA-specific cells within the GC B cell pool (**Fig. 2E**). When the total number was enumerated, OVA-specific IgG-switched B cells were around 5-fold (5.03-fold overall) more abundant in OVA/CFA-immunized non-Tg mice than act.OVA counterparts (**Fig. 3B**) and this was relatively similar to the relative abundance in the OVA-specific GC compartment (**Fig 2F**, 4.27-fold overall). These data strongly indicate that although fewer OVA-specific B cells appear to be recruited into GC reactions in act.OVA mice, once recruited they proceed through class-switch recombination (CSR) with equivalent efficiency as in non-Tg mice. To confirm whether this was the case we next probed for OVA-specific B cells that may have failed to undergo CSR in act.OVA mice. Looking within the unswitched OVA-specific B cell populations it was apparent that there was no major enrichment of unswitched OVA-specific B cells in act.OVA mice relative to non-Tg controls (**Supplementary Figure 2C,D**) and consistent with this virtually all OVA-specific GC B cells in OVA/CFA immunized non-Tg and act.OVA mice had undergone switching (**Supplementary Figure 2E**).

**Figure 3.**
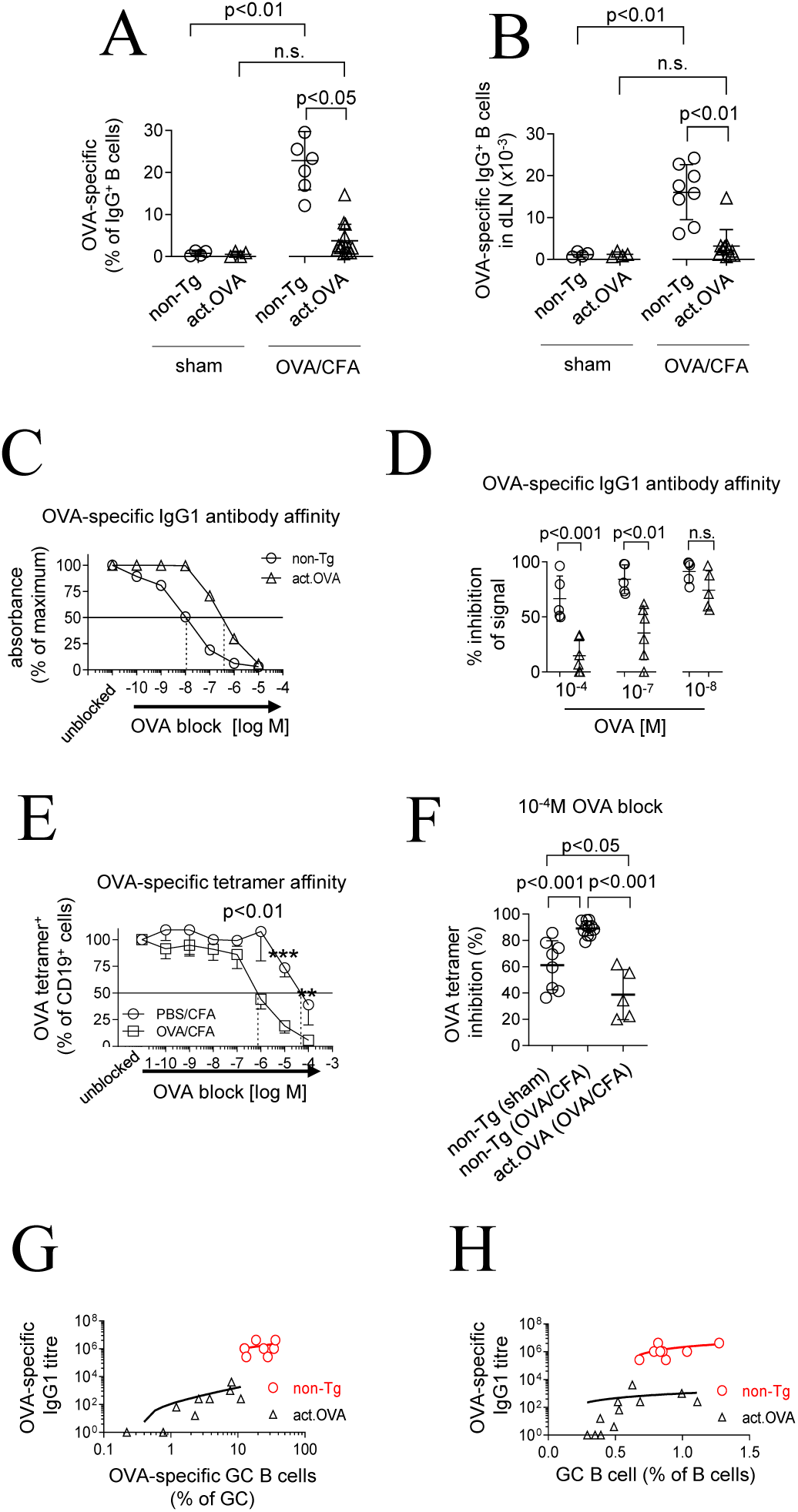
OVA-specific CSR and affinity maturation are reduced in act.OVA mice. Non-Tg or act.OVA mice were sham-immunized (PBS/CFA) or immunized with OVA/CFA and analysed 14 days later. A, B) Accumulation of OVA-specific IgG^+^ switched B cells in dLN. C, D) Relative affinity of OVA-specific IgG1 antibodies in plasma collected 14 days after immunization was compared using an ELISA-based assay. E) Blockade of tetramer binding by titrated concentrations of soluble OVA for B cells from dLN taken from sham- or OVA/CFA-immunized non-Tg mice shown as proportion (%) of OVA-tetramer^+^ cells relative to unblocked. F) Extent of tetramer binding inhibition by 10^−4^M OVA. G,H) Correlation of OVA-specific IgG1 titre and GC B cell number. Data show values for individual mice with mean±SD (A,B), absorbance (% of maximal) vs concentration of OVA block (C), % reduction in absorbance relative to unblocked samples at the indicated concentrations of OVA for individual mice pooled from 3 experiments with mean±SD (D), % reduction in OVA-tetramer^+^ cells for blocked relative to unblocked samples showing mean±SD (n=4/analysis point) (E) or individual mice showing mean±SD (F) and IgG1 titre vs OVA-specific (G) or total (H) GC B cell frequency for individual mice. Data are representative (C) or pooled from 2-5 experiments. Kruskal-Wallis (A,B), ANOVA with Tukey’s post-test (F), t-test (D,E), log/log (G) or log/linear regression (H).

To understand whether affinity maturation was modulated, we compared the relative affinity of circulating OVA-specific IgG1 from non-Tg controls and act.OVA mice after OVA/CFA immunization using an ELISA-based competitive blocking assay. Comparing the IC_50_, the affinity of OVA-specific IgG1 present in the circulation 14 days after immunization was reduced by approximately 2 orders of magnitude in act.OVA mice relative to non-Tg controls (**Fig. 3C**). This was most evident when comparing samples blocked with a molar concentration of OVA around the IC_50_ for OVA/CFA immunized non-Tg controls (10^−7^ – 10^−8^M; **Fig. 3D**). We next probed affinity directly at the cellular level using an assay we developed where preincubation with titrated concentrations of soluble OVA is used to block OVA tetramer binding to reveal differences in surface BCR affinity (17). Tetramer binding to B cells from dLN of OVA/CFA immunized non-Tg controls was inhibited by ∼100-fold lower concentrations of soluble OVA than for B cells from OVA/CFA immunized act.OVA mice indicating a ∼100-fold greater BCR affinity for the cells from non-Tg mice. Notably, 10^−4^M soluble OVA effectively blocked tetramer binding to the affinity-matured OVA-specific BCR in OVA/CFA immunized non-Tg mice but only partially blocked binding to the presumptively non-matured BCR in act.OVA mice (**Fig. 3E**). Using 10^−4^M soluble OVA block, there was less inhibition of tetramer binding for B cells from OVA/CFA immunized act.OVA mice than for sham or OVA/CFA immunized non-Tg mice (**Fig. 3F**) indicating the presence of low affinity BCR on OVA-specific B cells in OVA/CFA immunized act.OVA mice. The very low frequency of OVA-specific B cells in act.OVA mice precludes analysis of their relative BCR affinity. This data suggested that although OVA-specific B cells had contributed to GC reactions (e.g. **Fig. 2**), affinity was not increased across the OVA-specific B-cell population when OVA was ubiquitously expressed.

As might be predicted, the OVA-specific antibody output was related to the frequency of OVA-specific B cells within the total GC B cell pool in act.OVA and non-Tg mice (**Fig. 3G)** and to the total GC size (**Fig. 3H**), suggesting that CFA-derived stimuli contribute substantially to OVA-specific antibody production in act.OVA mice but, on a per-cell basis, GC B cells in act.OVA mice appeared hampered in their capacity to produce OVA-specific IgG1. We propose that the GC checkpoint leads to counterselection of OVA-specific B cells which might have initially gained entry to the GC reaction as a result of accessory signals from CFA. Therefore, not only is the production of OVA-specific antibodies curtailed, affinity maturation is substantially limited. Additionally, there was also no difference in surface IgG levels (**Supplementary Figure 2F**), suggesting that the reduction of OVA-specific cells in the GC in act.OVA mice was not due to unsuccessful competition for antigen consequent to attenuated receptor expression but rather because of low BCR affinity for OVA.

### Transient antibody production reflects failed plasma cell differentiation in act.OVA mice

Our data thus far indicate that OVA-specific B-cell activation, and progression through GC differentiation and affinity maturation is substantially blunted, but OVA-specific GC and IgG-switched B cells do in fact accumulate, albeit at significantly reduced levels, after OVA/CFA immunization of act.OVA mice. Additionally, the antibody kinetics in act.OVA mice is consistent with a reduced output of OVA-specific plasma cells from GC reactions. Therefore, to gain insight into whether the low levels of OVA-specific IgG that accumulates might reflects reduced OVA-specific GC activity in act.OVA mice, we next examined plasmablast and plasma cell formation.

Globally, the proportion (**Supplementary Figure 3B,C**) and total number (**Fig. 4A-C**) of plasmablasts and plasma cells were similar between act.OVA and non-Tg mice after OVA/CFA immunization. Plasmablasts, which retain surface Ig expression, could be directly surveyed by OVA tetramer (**Fig. 4D**), and as expected, OVA-specific plasmablasts were almost completely absent from the plasmablast pool of OVA/CFA immunized act.OVA mice relative to non-Tg counterparts (**Fig. 4E,F**). To identify OVA-specific plasma cells, which do not retain surface Ig (**Supplementary Figure 3D**), we used intracellular OVA-tetramer staining. Validating this, OVA-specific plasma cells were present among lineage^-ve^CD19^var^CD138^+^ cells in the dLN following immunization (**Fig. 4G**), but not in the irrelevant CD138^-ve^lineage^+^ population (**Fig. 4G**). Specificity of detection was high as blocking with OVA prior to tetramer staining abolished tetramer binding to plasma cells (**Fig. 4G**). When we applied this analysis method to OVA/CFA-immunized act.OVA mice, we found virtually no OVA-specific plasma cells within the plasma cell pool in dLN (**Fig. 4H,I**) and, significantly, in bone marrow (**Fig. 4J**), the expected niche for long-lived plasma cells. In contrast, OVA-specific plasma cells were readily detected in OVA/CFA immunized non-Tg controls (**Fig. 4I,J**). Together, these data demonstrate that co-operation between GC and plasma cell differentiation checkpoints limit the lifespan and affinity of OVA-specific antibody production.

**Figure 4.**
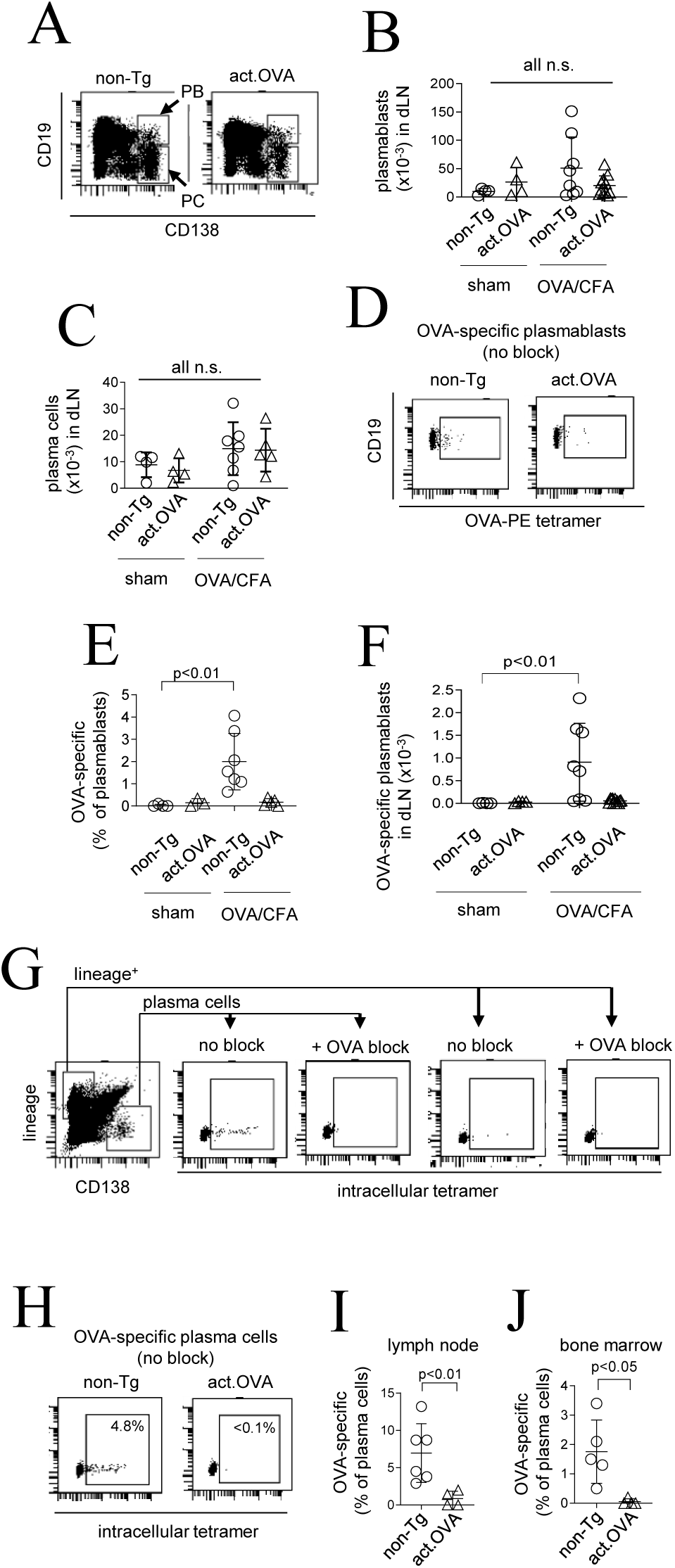
OVA-specific plasma cell responses are blunted in act.OVA mice. Non-Tg and act.OVA mice were sham-immunized (PBS/CFA) or immunized with OVA/CFA and dLN (pooled inguinal LN) and bone marrow was analysed 14 days later. A) Representative gating for plasmablasts (PB) and plasma cells (PC) based on CD19 and CD138 expression within lin^-^/CD19^var^ cells as shown in Suppl Fig 1. B) Total plasmablast number in dLN. C) Total number of plasma cells in dLN. D) Representative surface OVA-tetramer staining of plasmablasts in dLN with no OVA blocking. E) Proportion and total number of OVA-specific plasmablasts within all plasmablasts in dLN. F) Total number of OVA-specific plasmablasts in dLN G) Representative intracellular OVA tetramer staining of the indicated populations with and without OVA block (gated on CD19^+^ cells). H) Intracellular OVA-tetramer staining of plasma cells in dLN from non-Tg and act.OVA mice with no OVA blocking. (I,J) Frequency of OVA-specific plasma cells in dLN and bone marrow. Data show representative FACS plots (A,D,G,H) or individual mice with mean±SD (B,C,E,F,I,J). Data are pooled from 2-3 experiments. ANOVA with Tukey’s post-test (B,C,E), Kruskal-Wallis test (F), Mann-Whitney test (I,J).

### OVA-specific memory B cells fail to develop in OVA/CFA immunized act.OVA mice

Reactivation of Bmem mediates rapid antibody production upon antigen re-encounter. Based on our data thus far, we proposed that differentiation of OVA-specific memory B cells would be impaired in act.OVA mice. To determine whether OVA-specific Bmem arose as a consequence of OVA/CFA immunization we monitored the kinetics of Bmem formation (**Supplementary Figure 4A-E**) and then analysed dLN in detail 35 days post-immunization. We chose this timepoint because, among IgG-switched B cells in OVA/CFA immunized non-Tg and act.OVA mice, the Bmem pool remained relatively stable from this timepoint onwards but the abundance of GC B cells had waned from their peak level at day 14 (**Supplementary Figure 4C**). Additionally, among IgG-switched B cells, the Bmem compartment was dominant over the GC B cell compartment at day 35 (**Fig 5A, Supplementary Figure 4C-E**). Although the overall Bmem compartments were relatively similar between non-Tg controls and act.OVA mice (Supplementary Figure 4C,D), OVA-specific Bmem were remarkably rare in dLN of OVA/CFA-immunized act.OVA mice, but not non-Tg controls (**Fig. 5B-D**). Although there had been consistent but blunted development of GC B cells in dLN of OVA/CFA-immunized act.OVA mice 14 days after immunization (**Fig. 2**), at 35 days after immunization OVA-specific GC B cells were virtually absent from dLN (**Fig. 5E-G**) and considerably less abundant relative to non-Tg counterparts than at day 14 (72-fold less abundant at d35 compared to 4.3-fold less at day 14; **Fig. 2, Fig. 5G**). This suggests that the OVA-specific GC reaction in act.OVA mice diminishes much more quickly than in non-Tg controls, possibly as a result of limited OVA-specific B-cell survival as a consequence of limited T-cell help or through counterselection. Consistent with this there is virtually no circulating OVA-specific IgG1 detectable at this timepoint (**Fig. 1D, Supplementary Figure 4F**) indicating PB and PC differentiation has ceased in act.OVA mice.

**Figure 5.**
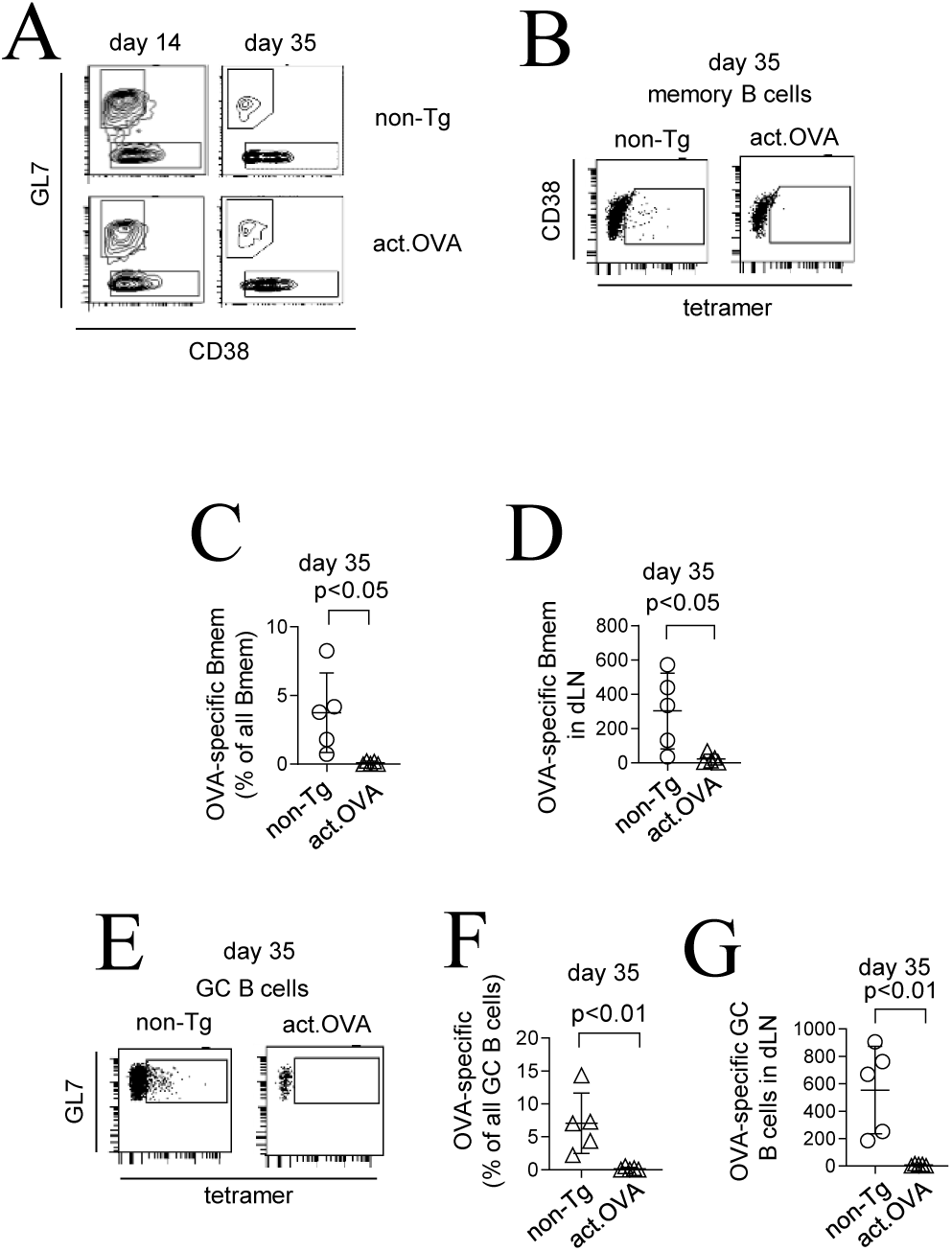
OVA-specific memory B cells fail to develop in act.OVA mice after immunization with immunogenic adjuvant. Non-Tg and act.OVA mice were immunized with OVA/CFA and lymph nodes were analysed at the indicated timepoints. A) Representative FACS plots showing GC (GL7+CD38-) and memory (CD38^+^GL7^-^) B cells, gated as per Supplementary Figure 1. B) Representative FACS plots showing OVA-tetramer staining of memory B cells at day 35, gated as per Supplementary Fig 1C) OVA-specific memory B cells as a proportion of all memory B cells in dLN at day 35. Total OVA-specific memory B cells in dLN at day 35. E) Representative FACS plots showing tetramer staining of GC B cells at day 35, gated as per Supplementary Fig 1. F) OVA-specific GC B cells as a proportion of all GC B cells at day 35. G) Total OVA-specific GC B cells in dLN at day 35. Data show representative FACS plots (A,B,E) or mean±SD or individual mice with mean±SD (C,D,F,G). Data are pooled from 2 experiments with 2-3 mice per group. Unpaired t-test (C,D,F,G).

Together these data confirm that when endogenous self-antigen is widely expressed, a GC checkpoint controls antigen-specific GC B cell survival and differentiation fate and efficiently terminates OVA-specific Bmem and plasma cell differentiation that arise as a consequence of bystander activation.

## Discussion

Many mechanisms combine to control development of unwanted, self-reactive B-cell responses and subsequent production of pathogenic autoantibodies. That naïve self-reactive B cells are relatively abundant in healthy individuals (21, 22) indicates the effectiveness of these mechanisms (12). Characterising how these mechanisms might fail is therefore critical to understanding the genesis of antibody-mediated autoimmune disease, and conversely, how tolerance approaches might be best deployed to achieve therapeutic goals. For convenience, studies of B-cell tolerance and its failure are typically performed in conditions where the BCR repertoire is genetically engineered to increase the frequency of the antigen-specific B cells to be studied, but this can alter tolerance outcomes. To understand the effectiveness of peripheral B-cell control mechanisms we studied autoantigen-specific B-cell responses, in a non-BCR transgenic repertoire, using adjuvants that provide bystander help through innate and T cell input. We found that only a highly potent adjuvant elicited antigen-specific autoantibody production and, even under these conditions, antibody production was transient. We also found that although CSR occurred, affinity maturation was blocked, and a post-GC checkpoint appeared to contribute to limiting antigen-specific antibody production.

Based on studies of BCR-transgenic mice, self-reactive B cells arriving in the periphery have effectively avoided deletion and/or receptor editing has insufficiently quenched the capacity to bind self. This represents only a minor fraction of the peripheral B cell pool, but residual self-reactivity must be contained by additional checkpoints. The stringency of these additional checkpoints has been tested using environmental perturbations (microbiota, infection) or genetic polymorphisms (autoimmune susceptibility genes). Even in non-autoimmune prone genetic backgrounds, some aspects of deletion, developmental arrest and anergy may be overcome by augmenting anti-apoptotic pathways (23) or by provision of T cell help (24) and innate stimuli (25). While these have been informative, the typical readouts for these experiments are technically limited and include the tracking of BCR-transgenic cells and/or indirect probing for the frequency of self-reactive BCRs by cloning and screening them as recombinant antibodies by ELISA (26). Whether these readouts directly reflect ongoing autoreactivity by a physiological repertoire remains unclear. Here we provide two important advances. First, we use sensitive and specific tetramer labelling to track the activation and progression of rare but endogenous self-reactive B cells through the GC reaction. Second, we couple analyses of antibody production and affinity to antigen-specific plasma cell differentiation to characterise the fate of GC-derived autoreactive B cells. Overall, our data are consistent with the well-appreciated role of tight regulation of GC entry and ongoing participation but highlight a novel checkpoint post-GC that limits long-lived antibody production.

Chronic sensing of self-antigen in the periphery enforces an anergy transcriptional program (10, 27) and the downregulation of surface IgM (26). The two key goals of this are 1) dampening signal transduction via the BCR upon self-ligation and 2) reducing the probability of encountering T cell help. Because GC entry is governed by T cell-B cell interaction, anergic B cells are anatomically alienated from follicles (4) and fail to receive necessary stimulation through surface co-receptors that enhance BCR signalling (3, 5, 24). One unresolved aspect is to what extent inflammation and non-cognate T cell help can drive promiscuous recruitment of autoreactive B cells into a GC reaction. It is well-documented that viral infection can lead to collateral autoantibodies (28) and, based on analyses of somatic hypermutation, that this can proceed through a canonical GC (29). Additionally, antibodies that are generated against pathogens are frequently polyreactive and cross-react with endogenous self-antigens (21). An oft-picked example is that antibodies neutralizing HIV are derived from germline autoreactive precursors (29), indicating that self-reactive B cell recruitment into a GC is a rare but biologically meaningful event. In a recent study (30), infection with γ-herpesvirus elicited affinity-matured autoantibodies that did not have germline autoreactivity, leading to the inference that self-binding had been acquired and positively selected inside GC. Our studies here using CFA agree in that total GC B cell frequency correlates with OVA-specific (auto)antibody output. We add further support to this by showing directly that self-reactive B cells can occupy GC following CFA immunization, albeit in a truncated manner, and that this is related to the magnitude of autoantibody production.

Survival and progression in the GC are driven in a Darwinian fashion by competition for limiting T cell help. Many if not all possible fates for GC B cells (cell cycle, deletion, memory and plasma cell differentiation) require T cell input (31). This is in part because GC B cell signalling is intrinsically rewired to depend on T cell help (31). Because the OVA-specific T cell compartment is tolerant (15), we suggest that sharp bystander signals from Mtb in CFA promote early self-reactive GC B cell responses at the time of immunization that were terminated through either a failure to compete in ongoing GC or through negative selection, or both. It is likely that providing an artificial source of cognate T cell help, either by adoptive transfer of primed OVA-specific CD4+ T cells or by covalently linking OVA to a foreign antigen, as others have done (32), autoreactive GC B cell responses may be either boosted or even rescued from deletion. Perhaps more useful, genetically enhancing positive signalling strength by modulating calcium flux (33) or removing negative regulation by inactivating inhibitory phosphatases (1) would reveal B cell-intrinsic contributions in addition to bystander T cell help, as shown here, that would be sufficient to drive bona fide autoimmunity. This would be particularly insightful where autoimmune susceptibility genes synergise with infection to initiate autoantibody production (34).

Because autoreactive GC B cells were not present at later timepoints and had not differentiated to memory B cells or long-lived plasma cells, we conclude that the GC shunted them towards deletion. This aligns with the low affinity of OVA-specific autoantibodies that were transiently produced, and that short-lived extrafollicular responses likely accounted for the transient autoantibody response. The rapid but transient kinetics and low affinity reported here resemble autoantibodies described from extrafollicular reactions that are relatively T-independent but for which innate signals (e.g microbe-derived LPS) can provide sufficient impetus for development (35). Indeed, innate signals, particularly those through TLR7 and TLR9 are sufficient to promote extrafollicular B-cell responses and autoantibody production (36-38) as well as T cell-independent CSR (39). Extrafollicular B cell activation and plasmablast differentiation may be a key site of autoantibody production, at least in RA (40, 41) and SLE (42), driven by self-antigens that are complexed with local TLR ligands like DNA or RNA (37, 38). We propose that in our studies, various Mtb-associated TLR signals (predominantly TLR2/4/9) serve a similar function and promote activation of peripheral anergic OVA-specific B cells, transiently breaking their state of functional unresponsiveness that is apparent in the absence of Mtb when IFA is used as adjuvant. Interestingly, although extrafollicular somatic hypermutation (40) can be driven by bacteria-associated (35, 41) or other TLR signals, it was not apparent here. This is most likely due to the way self-antigen is encountered by B cells. While the majority of studies characterising extrafollicular reactions are based on soluble self-antigen (40), the same membrane-bound self-antigens such as those employed here are more tolerogenic (6, 9) and provide more stringent control particularly when ubiquitously expressed in and around the B cell follicle. In any case, both GC and extrafollicular checkpoints operated to limit the production of self-reactive antibodies when tolerance is transiently perturbed.

The discrete stimuli that license self-reactive B cells is largely unknown in a polyclonal setting. These have instead been deduced using BCR-tg models which are useful tools to track defined populations of antigen-specific cells, however, qualitative and quantitative variations in tolerance outcomes have been reported when the BCR repertoire is genetically engineered. For example, the frequency of ‘engineered’ B cells is one determinant of outcomes as titrating in increasing numbers of alternative specificity ‘competitor’ B cells to increase ‘competition’ for survival factors like BAFF reduces self-reactive B cell lifespan (4, 43). In general, the frequency of engineered autoreactive B cells used in these studies exceeds their expected frequency in the endogenous repertoire by many orders of magnitude. In addition to the BCR-Tg B-cell frequency, the BCR-Tg model itself, the nature of the autoantigen, the BCR affinity and the genetic background, can modify sensitivity to signals controlling B-cell tolerance or responsiveness. As examples, LPS alone is not mitogenic for insulin-specific VH125xVk125 BCR-Tg B cells but is for other BCR-Tg B cells (e.g. 3H9 B cells) and upregulation of CD86 in response to LPS stimulation differs (1, 44, 45). For BCR-Tg mice generated with the VH10 HEL-specific BCR inserted ectopically into the genome, anergic B cells from HEL-Tg x soluble HEL-expressing mice are insensitive to LPS (46), whereas if the BCR construct is instead inserted into the endogenous IgH locus, B cells appear to show full responsiveness to LPS (47). Therefore, despite important insights gained from various BCR-Tg models and their experimental power for tracking antigen-specific B cells, there are some uncertainties remaining about how peripheral B-cell responsiveness is controlled, in a non-BCR engineered setting. We avoided such effects by analysing a non-BCR engineered setting. We show that in this setting that mechanisms proposed from studies of BCR-engineered settings largely hold true.

The findings here provide insight into not only the control but also the development of autoantibodies. They implicate a mechanism by which switched (low–affinity) self-reactive antibodies may arise in a polyclonal repertoire, when co-infection occurs in the presence of locally released self-antigens. Our findings show that some mechanisms defined in BCR-Tg models hold here in this setting that reflects a normal physiological polyclonal repertoire and refine our understanding of the control of B-cell responses. We propose our studies, performed here in a non-autoimmune-prone setting, along with tools we have developed (17) will provide valuable insights into understanding and controlling autoimmunity but also forms the basis of a valuable system for studying genetic susceptibility alleles or the role of individual molecules in peripheral control of B-cell responses. Our data which imply a critical role for T-cell help indicate that therapeutic approaches that directly modulate B-cells and T-cells and that counter genetic susceptibilities, such as HSC-based gene therapy (48) are likely to be highly effective at modulating autoantibody and other unwanted production.

## Acknowledgments

This research was carried out at the Translational Research Institute, Woolloongabba, QLD 4102, Australia. The Translational Research Institute is supported by a grant from the Australian Government. We acknowledge the Translational Research Institute core facilities, in particular the Flow Cytometry Core for their outstanding technical support.

## SUPPLEMENTARY FIGURES

**Supplementary Figure 1.**
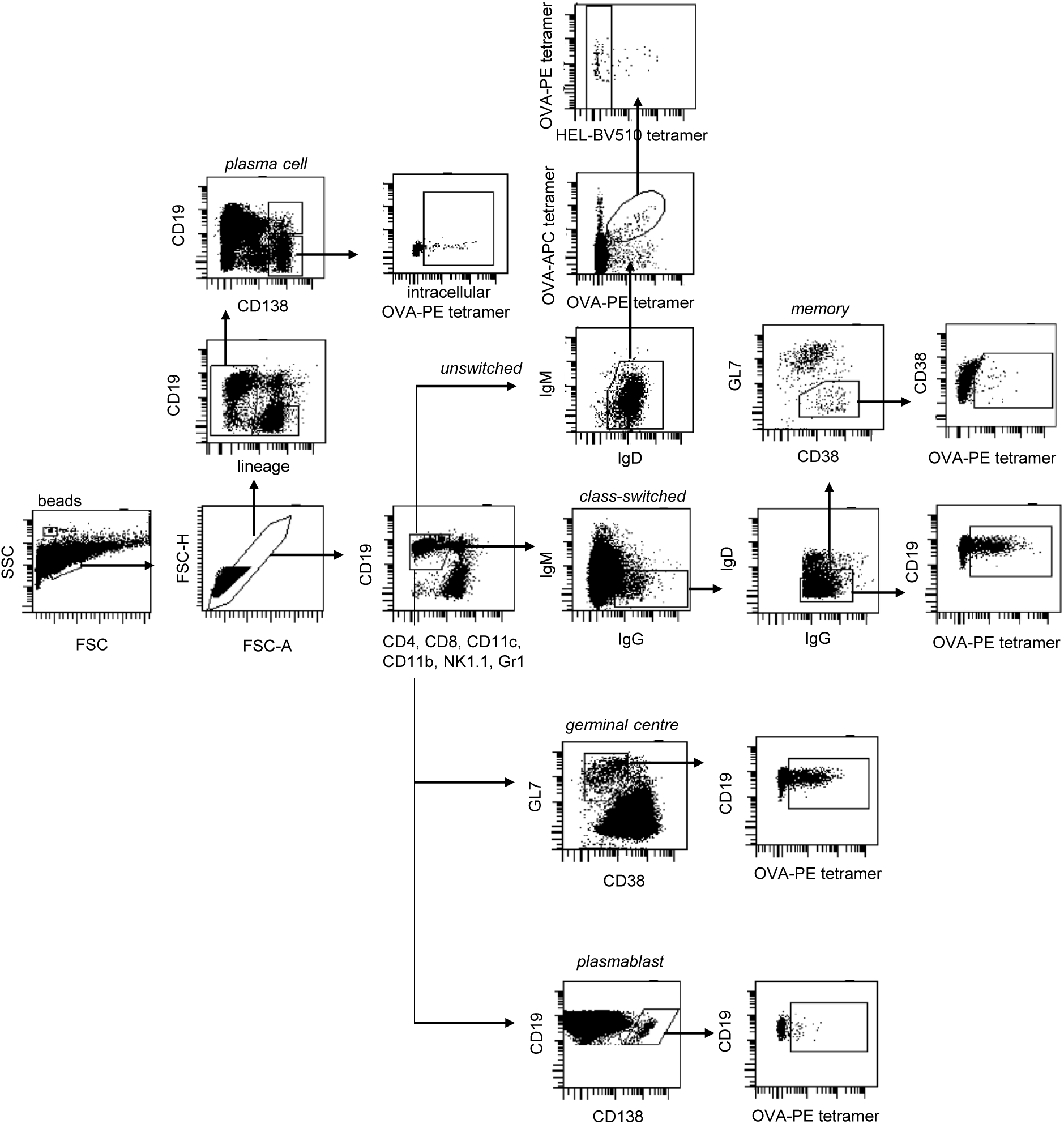
Representative gating strategy for B cell subsets, plasmablasts and plasma cells. FACS plots showing indicative staining of draining (pooled inguinal) lymph nodes from non-transgenic mice following immunization with OVA/CFA. Figure is a compilation to include subsets that were not necessarily analysed together for each single mouse. Memory B cell plots are from day 35 post-immunization, whereas all other plots are from day 14 post-immunization.

**Supplementary Figure 2.**
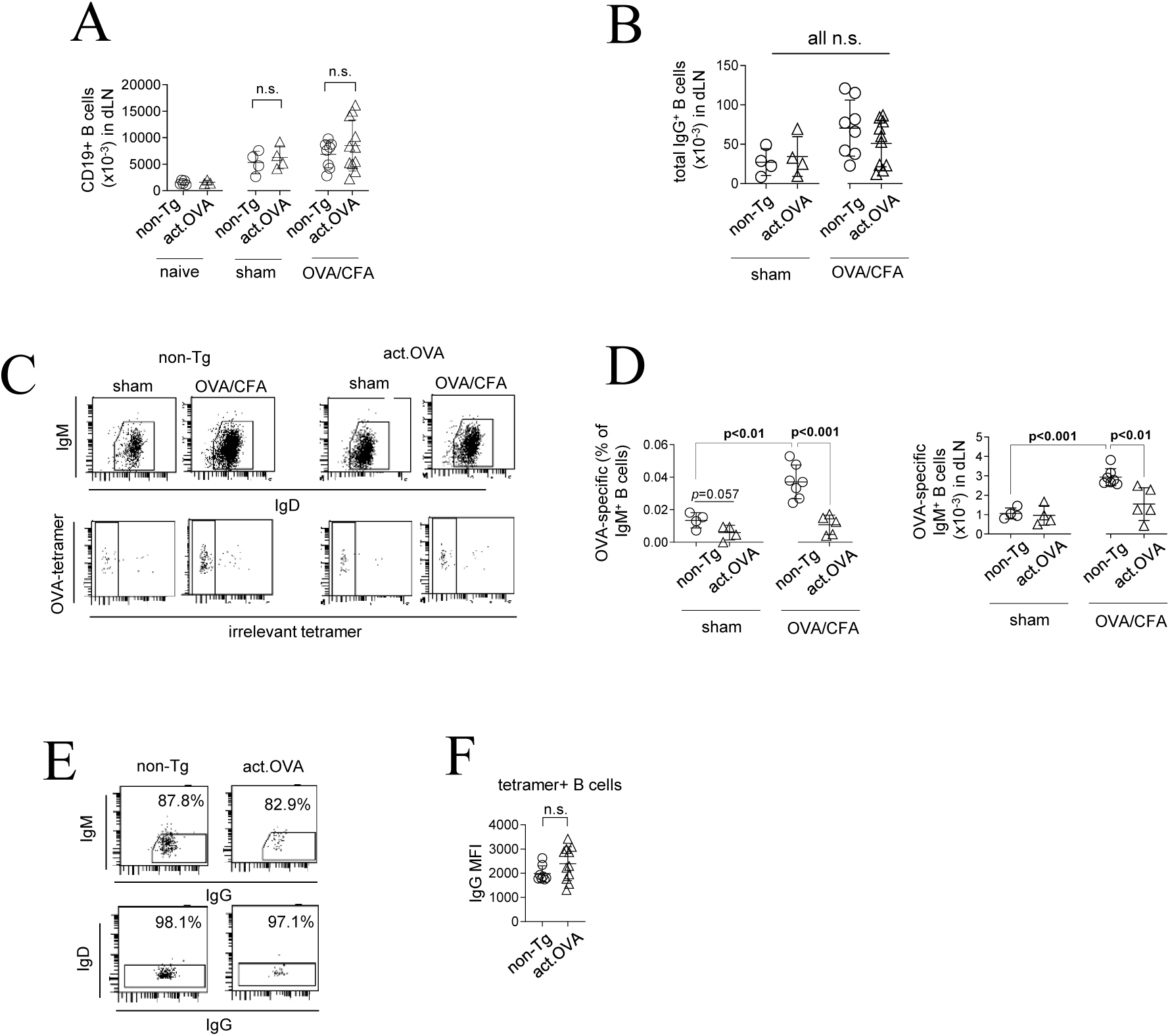
Analysis of switched and unswitched OVA-specific B cells in draining lymph node after OVA/CFA immunization. Act.OVA mice and non-transgenic littermates were left unimmunized (naïve), sham-immunized (PBS/CFA) or immunized with OVA/CFA and inguinal lymph nodes (dLN) were analysed 14 days later. A) Total number of CD19^+^ B cells in LN. B) Total number of IgG-switched B cells in dLN. C) Representative FACS plots showing unswitched B cells within the lin^-^CD19^+^ population (top panel) and OVA-specific B cells (negative for irrelevant tetramer) (bottom panel) gated as per Supplementary Figure 1. D) OVA-specific unswitched B cells as a proportion of all CD19^+^ cells in dLN (left) or total number in dLN (right). E) Representative FACS plots gated for OVA-specific germinal centre B cells. F) IgG MFI on OVA-specific B cells in the GC. Data show representative FACS plots (C, E) or values for individual mice with mean±SD. ANOVA with Tukey’s post-test (A, B, D) or unpaired t-test (F).

**Supplementary Figure 3.**
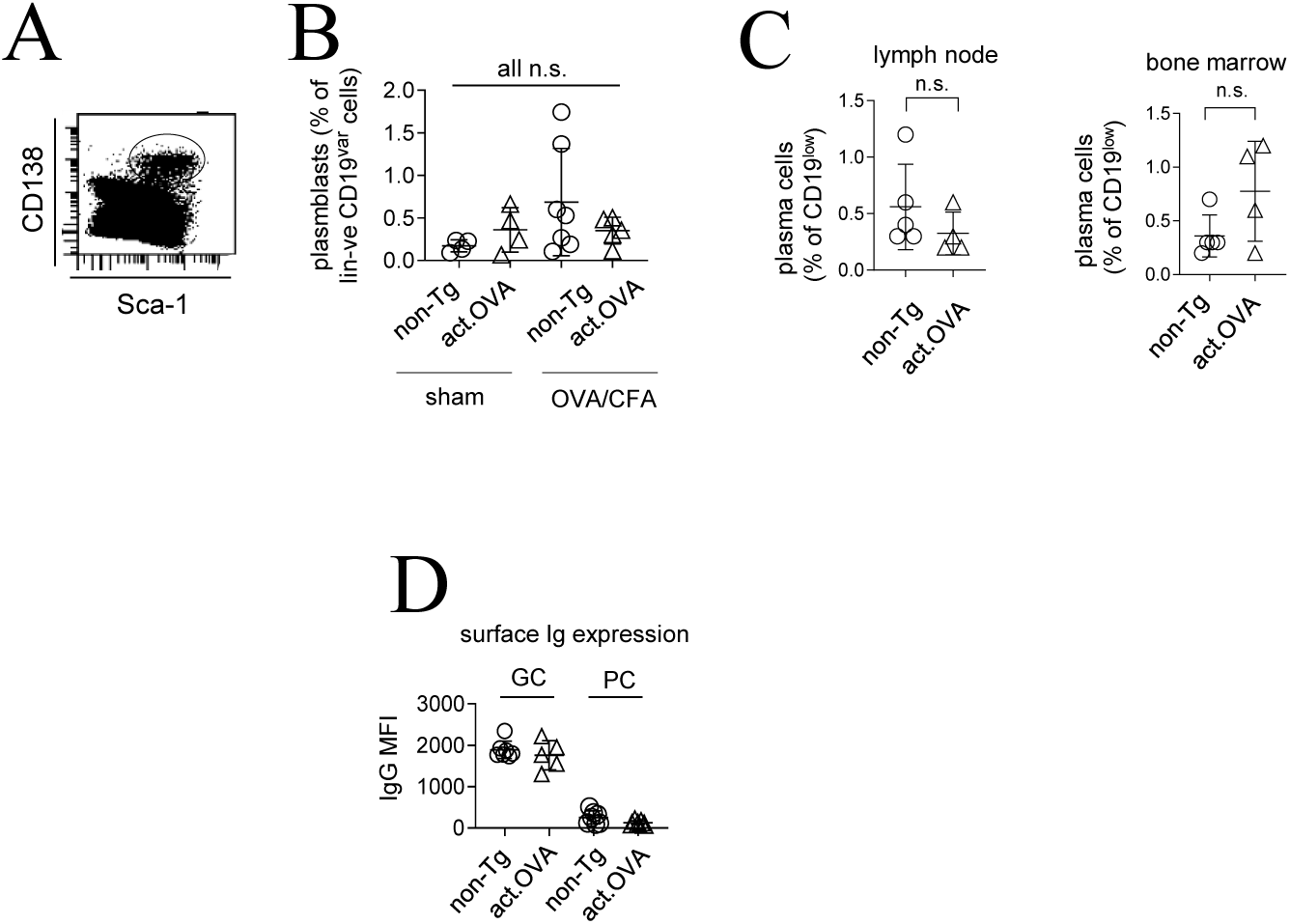
Differentiation of plasmablasts and plasma cells following immunization. Act.OVA mice and non-transgenic littermates were immunized with OVA/CFA and lymph nodes analysed 14 days later. A) Representative FACS plot of lin^-^CD19^var^ cells from non-transgenic lymph node, gated here for CD138 and Sca-1 expression. This staining was used to confirm that CD138 was marking plasma cells (Wilmore et al., Eur J Immunol (2017) 47:1386-1388). B) Proportion of plasmablasts (CD138^+^CD19^+^) within all B cells in dLN. Total plasma cells as a proportion of all CD19^var^ cells in lymph node or bone marrow. Surface IgG expression on germinal centre (GC) B cells and plasma cells (PC). This plot demonstrates that plasma cells cannot be directly detected by surface tetramer staining due to low/absent surface Ig expression. Data show representative FACS plot (A) or values for individual mice with mean±SD. ANOVA with Tukey’s post-test (B), t-test (C).

**Supplementary Figure 4.**
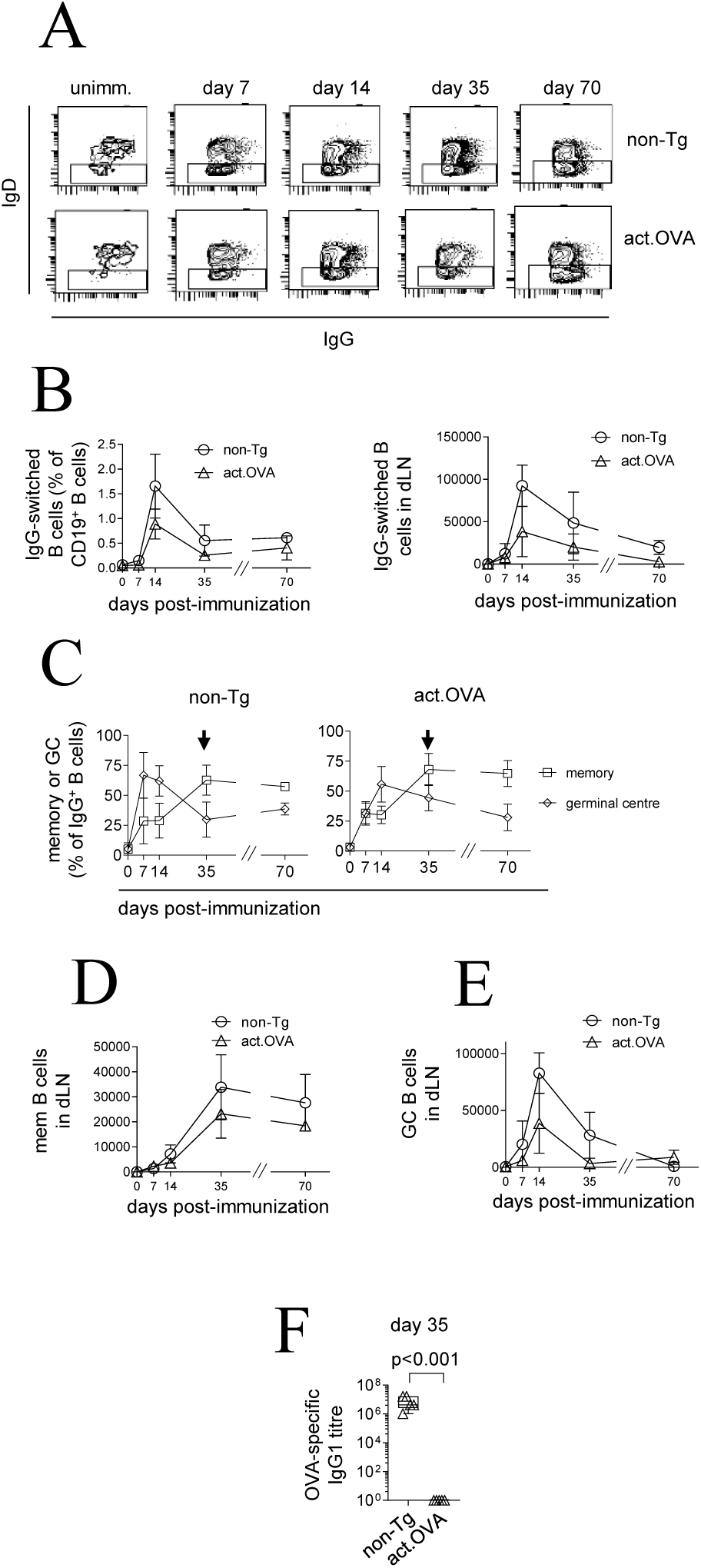
OVA-specific memory cells do not develop following immunogenic priming. Act.OVA mice and non-transgenic littermates were immunized with OVA/CFA and lymph nodes or plasma analysed at various timepoints. A) Representative FACS plots of IgG^+^ cells negative for IgM and IgD, pre-gated on IgM-cells as per Supplementary Figure 1. B) IgG^+^ B cells as a proportion of all B cells (right) or total number of IgG^+^ B cells (left) sampled at various timepoints. C) Proportion of memory or GC B cells sampled at various timepoints. D) Total memory B cells sampled at various timepoints. E) Total GC B cells sampled at various timepoints. F) OVA-specific IgG1 titre in plasma at day 35. Data show representative FACS plots (A), mean±SD (B-E) or values for individual mice showing mean±SD (F). t-test of log-transfomed titre (F).

